# Genome-wide association and prediction study in grapevine deciphers the genetic architecture of multiple traits and identifies genes under many new QTLs

**DOI:** 10.1101/2020.09.10.290890

**Authors:** Timothée Flutre, Loïc Le Cunff, Agota Fodor, Amandine Launay, Charles Romieu, Gilles Berger, Yves Bertrand, Nancy Terrier, Isabelle Beccavin, Virginie Bouckenooghe, Maryline Roques, Lucie Pinasseau, Arnaud Verbaere, Nicolas Sommerer, Véronique Cheynier, Roberto Bacilieri, Jean-Michel Boursiquot, Thierry Lacombe, Valérie Laucou, Patrice This, Jean-Pierre Péros, Agnès Doligez

## Abstract

To cope with the challenges faced by agriculture, speeding-up breeding programs is a worthy endeavor, especially for perennials such as grapevine, but requires understanding the genetic architecture of target traits. To go beyond the mapping of quantitative trait locus (QTL) in bi-parental crosses, we exploited a diverse panel of 279 *Vitis vinifera* L. cultivars. This panel planted in five blocks in the vineyard was phenotyped over several years for 127 traits including yield components, organic acids, aroma precursors, polyphenols, and a water stress indicator. The panel was genotyped for 63k single nucleotide polymorphisms (SNPs) by combining an 18K microarray and genotyping-by-sequencing (GBS). The experimental design allowed to reliably assess the genotypic values for most traits. Marker densification via GBS markedly increased the proportion of genetic variance explained by SNPs, and two multi-SNP models identified QTLs not found by a SNP-by-SNP model. Overall, 489 reliable QTLs were detected for 41% more response variables than by a SNP-by-SNP model with microarray-only SNPs, many new ones compared to the results from bi-parental crosses. Prediction accuracy higher than 0.42 was obtained for 50% of the response variables. Our overall approach as well as QTL and prediction results provide insights into the genetic architecture of target traits. New candidate genes and the application in breeding are discussed.

## Introduction

With the two major challenges facing perennial fruit crops in general and grapevine in particular, decreasing phytosanitary products such as fungicide treatments and adapting to climate change, both harnessing existing genetic diversity (Wolkovich et al. 2018) and breeding new varieties (Adam-Blondon et al. 2011) are important levers. For the latter, many studies aimed at deciphering the genetic architecture of traits of interest by mapping quantitative trait locus (QTL) in bi-parental progenies (Vezzulli et al. 2019). However, this approach suffers from several drawbacks: the limited allelic diversity in parents, the low number of recombination events in the progeny, the upward bias of estimated QTL effects, and the under-estimation of the polygenic contribution for prediction purposes (Cardon and Bell 2001). As a result, all traits currently involved in grapevine marker-assisted selection (Vezzulli et al. 2019) are controlled by major genes, such as resistance to downy and powdery mildews (Di Gaspero et al. 2007), black rot (Rex et al. 2014), sex (Picq et al. 2014), berry color (Fournier-Level et al. 2009), seedlessness (Mejía et al. 2011), and Muscat aroma (Duchêne et al. 2009).

To overcome these limits, a few genome-wide association studies (GWASs) were performed in cultivated grapevine diversity panels but did not identify many new QTLs, due to various reasons. Several articles (Myles et al. 2011; Zarouri 2016; Migicovsky et al. 2017; Laucou et al. 2018) harnessed phenotypic data from genetic resources repositories collected without a proper experimental design. Moreover, the first three cited articles used at most 10k SNPs despite the low extent of linkage disequilibrium (Myles et al. 2011; Nicolas et al. 2016). Among other articles, Zhang et al. 2017 focused on a single binary trait with a major QTL, seedlessness; Yang et al. 2017 used only 187 SSRs and 96 genotypes; Sargolzaei et al. 2020 focussed on disease resistance using a 18K SNP microarray; Naegele et al. 2021 used at most 14k SNPs obtained by sequencing.

Moreover, most of these articles as well as one with 32k SNPs obtained with sequencing (Guo et al. 2019) used SNP-by-SNP models. However, multi-SNP models have the advantage of explicitly assuming a genetic architecture, be it sparse with few major QTLs or dense with many small-effect QTLs, allowing them to benefit from a potential gain in power (Zhang et al. 2019). Furthermore, the effects of QTLs often are overestimated (Xu 2003) which leads to poor prediction (Meuwissen et al. 2001). Multi-SNP models provide a natural way to efficiently perform genomic prediction (de los Campos et al. 2013), notably for traits with no major QTLs.

Consequently, our objective was to perform whole-genome association and prediction for various traits of interest in grapevine breeding, likely to display different genetic architectures. We aimed at finding out to what extent genetic variation contributes to phenotypic variation, how it is organized in sparse and dense genetic components, how accurate genomic prediction might be, and which genes are present under the QTLs. Our approach builds on a large diversity panel of 279 *Vitis vinifera* L. cultivars (Nicolas et al. 2016) defined from the French collection of genetic resources and overgrafted in the vineyard in five randomized complete blocks. The panel was phenotyped over several years and under different conditions for 127 traits, including yield components, organic acids, aroma precursors, polyphenols, and a water stress indicator, which, along with 25 derived variables, totaled 152 response variables. The cultivars were genotyped with both microarray and sequencing after a reduction of genomic complexity (genotyping-by-sequencing, GBS), reaching a total of 63k SNPs. QTL detection and genomic prediction were then performed with multi-SNP models assuming different genetic architectures, and positional candidate genes were searched for under QTLs.

## Materials and methods

### Plant material and field trial

The panel of 279 cultivars of *Vitis vinifera* L. is weakly structured in three genetic groups (Nicolas et al. 2016). In 2009, at the Domaine du Chapitre of the Institut Agro (Villeneuve-lès-Maguelone, France), the 279 cultivars as well as a control (cv. Marselan) were all overgrafted on 6-year-old vines of Marselan, itself grafted on rootstock Fercal, in a complete randomized block design with five blocks (A to E, supplementary figure S1). Because of failed overgraft, precocious death or fertility issues, only 270 cultivars among the 279 from the whole panel could be phenotyped in the field. The density of the field trial was 3300 plants/ha (1 m between plants along the same rank and 2.5 m between ranks). Each of the five blocks contained one plant of each panel cultivar as well as a regular mesh of over-grafts of Marselan as control (between 23 and 39 per block). The double-cordon training system was applied.

A random subset of 21 full-sib genotypes of a Syrah x Grenache progeny, together with the 2 parents, was also used to assess out-of-sample genomic prediction using a field design already described (Doligez et al. 2013).

### Phenotyping

About terminology, we use the term “trait” for any plant feature for which raw data were collected, whatever the year and the condition. However, in our analyzes we use the term “response variable” because: (i) for some traits, data were acquired in different years and conditions, hence analyzed separately; (ii) we also combined several traits to define new variables. In the end, we hence analyzed 152 response variables from 127 traits.

In 2011 and 2012, the trial was not irrigated, and both the panel cultivars and the control were phenotyped. For each plant, we recorded the nuber of clusters (NCBLU) and harvested three clusters at 20°Brix, which provided the sampling date (SAMPLDAY, in days since January 1). We measured mean cluster weight (MCW, in g), mean cluster length (MCL, in cm), mean cluster width (MCWI, in cm) and cluster compactness (CLUCOMP, on the OIV 204 scale from 1 to 9; OIV, 2009). One hundred berries randomly sampled from the middle of clusters were weighted, providing the mean berry weight (MBW, in g). In 2011-2012 and 2012-2013 winters, the number of woody shoots (NBWS) and pruning weight (PRUW, in kg) were measured for each plant. In 2011, the veraison date (onset of ripening, VER, in days since January 1) was also recorded. Two variables were computed from these traits: the veraison-maturity interval (VERMATU as SAMPLDAY - VER, in days), and plant vigour (VIG as PRUW / NBWS, in kg). In 2011 and 2012, juices were made from the sampled berries and analyzed to measure δ^13^C (D13C) as previously detailed (Pinasseau, Vallverdú-Queralt, et al. 2017). In 2012 were also measured glucose (GLU), fructose (FRU), malate (MAL), tartrate (TAR), shikimate (SHI) and citrate (CIT) concentrations, all in μEq.L^-1^, as previously detailed (Rienth et al. 2016). Six variables were computed from these traits: the sum of glucose and fructose (GLUFRU), glucose divided by fructose (GLUONFRU), malate divided by tartrate (MALTAR), idem for shikimate (SHIKTAR), citrate (CITAR) and the sum of glucose and fructose (GLUFRUTAR).

In 2014 and 2015, irrigation was applied to blocks C, D and E only (Pinasseau, Vallverdú-Queralt, et al. 2017), and only panel cultivars were phenotyped. As above, three clusters per plant were harvested at 20°Brix, providing the mean cluster weight (MCW, in g). Berries sampled from different blocks with the same water treatment were pooled per cultivar. More details on berry sampling and processing, as well as polyphenols and δ^13^C measurements and analysis are described elsewhere (Pinasseau, Vallverdú-Queralt, et al. 2017). From the available data on the 105 polyphenols in μg per berry (Pinasseau, Verbaere, et al. 2017), a few typos were corrected and 17 extra variables were calculated (Pinasseau, Vallverdú-Queralt, et al. 2017). In addition, two aroma precursors, β-damascenone (BDAM, in μg.L^-1^) (Kotseridis et al. 1999) and potential dimethyl sulfide (PDMS, in μg.L^-1^) (Segurel et al. 2005) were also measured. The volume and weight of juice samples were recorded, allowing to assess their effects when included as co-factors in the statistical analyses.

A total of 127 traits were phenotyped, from which 25 extra variables were computed. Because irrigation was applied to some blocks only in 2014-2015, the few traits phenotyped both in 2011-2012 and in 2014-2015 were analyzed separately. Overall, 152 response variables were analyzed (supplementary tables S1 and S2).

The sanitary status of cultivars regarding the presence of five viruses (CNa, GLRaV1, GLRaV2, GLRaV3, GFkV) was assessed by ELISA from plants at INRAE Vassal (Marseillan, France). Flower sex (OIV 151) and berry skin color (OIV 225) of each panel cultivar were retrieved from (Laucou et al. 2018) and completed with the database of INRAE Vassal germplasm repository (https://bioweb.supagro.inra.fr/collections_vigne/Home.php?l=EN).

Berry weight was phenotyped on the Syrah x Grenache cross in 2005, 2006 and 2007 in the same way as on the panel (Doligez et al. 2013), with 8 clusters harvested per genotype and per block instead of 3.

### Genotyping

#### Data acquisition and analysis of microarray SNPs

The panel and the Syrah x Grenache progeny were genotyped with the GrapeReSeq 18k Vitis Illumina microarray (Laucou et al. 2018). Data processing (see supplementary text S1, figures S2 and S3, and table S2) resulted in 13,925 SNPs for 277 cultivars. After filtering on linkage disequilibrium (LD) above 0.9 and minor allele frequency (MAF) below 0.05, 10,530 SNPs remained (see supplementary figure S4), thereafter referred to as the “microarray-only” SNPs.

#### Data acquisition and analysis of sequencing SNPs

The panel was also genotyped by sequencing (GBS, Elshire et al. 2011). Keygene N.V. owns patents and patent applications protecting its Sequence Based Genotyping technologies. Data processing consisted in read checking with FastQC version 0.1.2 (Andrews 2016), demultiplexing with a custom script, cleaning and trimming with CutAdapt version 1.8.1 (Martin 2011), alignment on the PN40024 12Xv2 reference sequence (Canaguier et al. 2017) with BWA MEM version 0.7.12-r1039 (Li 2013 Mar 16) and re-alignment with GATK version 3.7 (DePristo et al. 2011), followed by variant and SNP calling with GATK HaplotypeCaller, and a final filtering step, notably to discard SNP genotypes with less than 10 reads or quality below 20 (see supplementary text S2 and table S3). It resulted in 184,145 SNPs with less than 30% missing data for the 279 panel cultivars.

#### Joint imputation of microarray and GBS SNPs

The 13,925 microarray SNPs and 184,145 GBS SNPs for 277 common cultivars were combined with duplicate removal into a set of 197,885 common SNPs using coordinates on the 12Xv2 reference sequence (Canaguier et al. 2017). Missing data were imputed using LD with Beagle version 4.1-r862 (Browning and Browning 2009) as advised by Swarts et al. (2014), with window=1000, overlap=450, ne=10000 and otherwise default parameters. After filtering for LD above 0.9, 90,007 SNPs remained (see supplementary figure S3), and after filtering for MAF below 0.05, 63,105 SNPs remained (see supplementary figure S4), thereafter referred to as the “microarray-GBS” SNPs. We also imputed the Syrah x Grenache SNP genotypes similarly using Beagle.

### Statistical modeling of phenotypic data

We performed a two-stage analysis of each response variable using univariate regression models. In the first stage, estimates of total genotypic values were obtained (detailed in this section). In the second stage (see next section), these were regressed on SNP genotypes to identify QTLs, estimate their allelic effects and assess prediction accuracy.

To decrease the influence of potential outliers, all polyphenols (the compounds as well as the calculated variables) had their raw data automatically transformed with the natural log. For the other traits, when their raw phenotypic data were too skewed as visually assessed, they were also transformed with the natural log (see supplementary figure S5 and table S4).

#### Assessment of spatial heterogeneity

In 2011-2012, phenotypic data for the control were spatially analyzed (Cressie 1993) in a way similar to (Hamann et al. 2002). First, a global linear model was fitted with R/stats with fixed effects for block, year, block-year interaction, pruning weight, number of woody shoots, vigour, and all five viruses (pruning weight and vigour were discarded when vigour itself was the response). Facilitated by R/MuMIn version 1.40 (Bartoń 2017), model comparison was performed by maximum likelihood (ML), the best model being selected based on the corrected Akaike information criterion, AICc (Burnham and Anderson 2004). For each year separately, the empirical variogram of residuals from the best model was computed, on which several variogram models were fitted by ML with R/gstat version 1.1.5 (Pebesma 2004): exponential, spherical, gaussian and Stein’s parametrization of the Matérn model. The variogram model with the smallest sum of squared errors was then used to perform spatial interpolation by kriging, i.e. best linear unbiased prediction (BLUP) of the control’s response variable at all locations. By visually assessing the slope of the best variogram model fitted to the empirical variogram (supplementary figure S6) and the prediction errors from cross-validation (not shown), it was concluded that there was no need to correct for spatial heterogeneity.

In 2014-2015, the control was not phenotyped, an irrigation treatment was applied, and samples from different blocks with the same irrigation level were pooled (Pinasseau, Vallverdú-Queralt, et al. 2017), hence preventing the assessment of potential spatial heterogeneity as above.

#### Estimation of genotypic values

For each response variable, a global linear mixed model was defined with multiple fixed effects (for the 2011-2012 data set: block, year, block-year interaction, pruning weight, number of woody shoots, vigour, and all five viruses, pruning weight and vigour being discarded when vigour itself was the response; for the 2014-2015 data set: irrigation, year, irrigation-year interaction, °Brix (as their can be small deviations from 20°Brix), and all five viruses, as well as the volume and weight of juice samples for BDAM and PDMS) together with two random effects (genotype and genotype-year interaction). The global model was fitted by maximum likelihood (ML) with R/lme4 version 1.1.19 (Bates et al. 2015). The output was given to R/lmerTest version 3.1-2 (Kuznetsova et al. 2017) to use its function “step.” Backward elimination of random-effect terms was performed using likelihood ratio test, followed by backward elimination of fixed-effect terms using F-test for all marginal terms, i.e., terms that can be dropped from the model while respecting the hierarchy of terms in the model, with a threshold on *p* values at 0.05 for both types of terms. The final model after backward elimination was then re-fitted by restricted maximum likelihood (ReML) to obtain unbiased estimates of variance components and empirical BLUPs of genotypic values. The acceptability of underlying assumptions (homoscedasticity, normality, independence) was visually assessed by plotting residuals and BLUPs. Broad-sense heritability on a genotype-mean basis (H^2^) was computed using two estimators. The first assumes a balanced design (Falconer and Mackay 2009): H^2^_C_ = σ^2^_g_ / [σ^2^_g_ + (σ^2^_g:y_ / n_y_) + (σ^2^_e_ / (n_y_ x n_r_)] where σ^2^_g_ is the variance of the genotypic values, σ^2^_g:y_ is the variance of the genotype-year interactions, n_y_ is the arithmetic mean number of trials (years), σ^2^_e_ is the variance of the errors (residuals) and n_r_ the arithmetic mean number of replicates per trial. The second estimator allows unbalanced data, H^2^_O_ (see Oakey et al., 2006, for details). Robust confidence intervals for variance components, heritability and genotypic coefficient of variation were obtained by parametric bootstrap as recommended by Schweiger et al. (2016), using the percentile method (Carpenter and Bithell 2000) in the R/lme4 and R/boot packages. In the Syrah x Grenache progeny, empirical BLUPs of genotypic values for berry weight were obtained in the same way.

### Statistical modeling of genotypic data

#### Genetic architecture assumed sparse

We used two types of models to perform genome-wide association testing and detect QTLs. The first is the SNP-by-SNP model as implemented in GEMMA version 0.97 (Zhou and Stephens 2012). For each SNP p, eBLUP(**g**) = **1** μ + M_a,p_ β_p_ + **u** + **e** where eBLUP(**g**) is a vector of responses of length N, M_a,p_ is a vector of length N with the genotypes at the p^th^ SNP (additive coding), β_p_ is its effect modeled as fixed, **u** ~ N_N_(**0**, σ_u_^2^ A) is a vector of length N corresponding to a polygenic effect modeled as random where the covariance matrix A contains additive genetic relationships (Vitezica et al. 2013), and **e** ~ N_N_(**0**, σ_e_^2^ Id) with N the Normal distribution of dimension N, **0** a vector of zeros and Id the identity matrix of dimension NxN. eBLUPs of **g** were used instead of BLUEs as they are known to be more accurate for prediction and selection purposes, notably thanks to the shrinkage property (Piepho et al. 2008). Our goal was to test the null hypothesis β_p_=0 while controlling for relatedness between genotypes. Controlling the family-wise error rate at 5% to account for multiple testing, the effect of a SNP was deemed significant when the *p* value from the Wald test statistic was lower than the Bonferroni threshold.

The second type of models jointly analyzes all SNPs with the goal of selecting a subset of them with large effects while handling linkage disequilibrium. This SNP selection can be achieved in a frequentist setting *via* stepwise regression (Segura et al. 2012). It starts with the SNP-by-SNP model, followed by inclusion, at every iteration, of the SNP with the smallest *p* value as an additional fixed effect, until the proportion of variance explained by the polygenic effect is close to zero. The SNP effects deemed significant were those of the best model selected according to the extended BIC. We fitted it with R/mlmm.gwas v1.0.4 (Bonnafous et al. 2018) allowing a maximum of 50 iterations. SNP selection can also be achieved in a Bayesian setting with the following model: eBLUP(**g**) = **1** μ + M_a_ **β** + **e**, where M_a_ is a NxP matrix of SNP genotypes (additive coding), with the so-called spike-and-slab prior for each SNP *p*, β_p_ ~ π_0_ δ_0_ + (1 - π_0_) N_1_(0, σ_β_^2^), δ_0_ being a point mass at zero. We fitted it with the variational algorithm, faster than MCMC, implemented in R/varbvs version 2.5.7 (Carbonetto and Stephens 2012). A SNP was deemed significant when its posterior inclusion probability, PIP_p_ = Pr(β_p_ ≠ 0), was larger than 0.80.

Beyond this focus on statistical significance (McShane and Gal 2017), we provide all estimates of significant additive SNP effects with a quantification of their uncertainty (supplementary table S5).

#### QTL definition and annotation

QTLs were defined as intervals around significant SNPs based on the decay of linkage disequilibrium (Bonnafous et al. 2018) (see supplementary text S3). A comparison was performed between the QTLs detected in this study and two lists, the first of already-published QTLs (Vezzulli et al. 2019), significant at a 5% genome-wide threshold, that were classified according to the Vitis INRAE ontology v2 (https://urgi.versailles.inra.fr/ephesis/ephesis) and slightly edited for automatic processing (see supplementary text S3), and the second of significant hits from a few GWAS publications after converting their coordinates on the genome reference we used.

In terms of annotation, as a given locus can be a QTL for multiple response variables, we first merged our 489 reliable QTLs (found with at least two methods, see Results) across all response variables, which resulted in 134 distinct genomic intervals (supplementary table S9). These intervals had a median length of 100,001 kb (with a minimum of 100,001 kb and a maximum of 1,072,169 kb). We then searched for overlaps between them and the Vcost version 3 annotations totalizing 42,413 gene models from (Canaguier et al. 2017), also using the correspondence between IGGP (International Grapevine Genome Program) and NCBI RefSeq gene model identifiers provided by the URGI (https://urgi.versailles.inra.fr/Species/Vitis/Annotations).

#### Genetic architecture assumed dense

To estimate the proportion of variance of empirical BLUPs of genotypic values explained by the cumulative contribution of SNPs (Yang et al. 2010) (PVE_SNPs_), we used the well-known multi-SNP model called ridge regression (also known as “RRBLUP”) which assumes a dense architecture: eBLUP(**g**) = **1** μ + M_a_ **β** + **e** where **β** ~ N_P_(**0**, σ_β_^2^ Id). It is known to be equivalent to the “GBLUP” model (Habier et al. 2007; Vitezica et al. 2013): eBLUP(**g**) = **1** μ + **g_a_** + **e** where **g_a_** ~ N_N_(**0**, σ_a_^2^ A) with A, the NxN matrix of additive genetic relationships, proportional to the matrix product M_a_ M_a_^T^ once M_a_ is centered using allele frequencies. Similarly for the dominance genotypic values **g_d_** ~ N_N_(**0**, σ_d_^2^ D) where D is the NxN matrix of dominance genetic relationships. Because the estimators of additive and dominance relationships from SNPs assume linkage equilibrium, a threshold on LD of 0.5 was applied when computing A and D. We fitted the models with R/lme4 and computed confidence intervals for variance components by bootstrap as above.

#### Genomic prediction

Out-of-sample prediction was assessed within the panel by 5-fold cross-validation repeated 10 times with R/caret version 6 (Kuhn 2018), using R/varbvs which assumes a sparse genetic architecture and R/rrBLUP version 4.5 (Endelman 2011) which assumes a dense architecture (infinitesimal model). Note that the QTL results from the GWAS analysis were not used when training each model, to avoid overfitting. We assessed prediction accuracy between empirical BLUPs of genotypic values and their predictions with various metrics: root mean square error (RMSE), Pearson’s linear correlation coefficient (corP), Spearman’s rank correlation coefficient (corS), as well as outputs from the simple linear regression of observations on predictions such as the intercept, slope, adjusted coefficient of determination (R^2^) and the *p* value of the test for no bias. Out-of-sample prediction was also assessed by training rrBLUP and varbvs methods on the whole panel and predicting empirical BLUPs of genotypic values for the 23 genotypes of the Syrah x Grenache cross.

## Results

### Estimation of broad-sense heritability and genetic coefficient of variation

We took advantage of the *Vitis vinifera* L. panel of 279 cultivars suitable for GWAS and representing the INRAE Vassal germplasm repository to set up a randomized-complete-block field trial (supplementary figure S1). It was phenotyped for 127 traits from which 25 extra variables were computed. All 152 response variables displayed substantial variation (supplementary figure S5). For some polyphenol variables, part of the variation was obviously associated with skin color, 137 cultivars out of 279 having colored skin berries. When phenotyped, the control cultivar allowed us to assess that (i) part of this variation is due to genetic differences between panel cultivars (supplementary figure S5), and that (ii) spatial heterogeneity is negligible (supplementary figure S6). The amount of missing data among response variables ranged from 15.78% to 43.93% (supplementary table S4). To account for such unbalance, we fitted linear mixed models and obtained BLUPs of genotypic values. After model selection, the final set of fixed and random effects differed between response variables (supplementary table S4), with year and genotype-year interaction effects being selected in most cases.

We then assessed the accuracy with which genotypic values were estimated using broadsense heritability (the higher, the better). As shown in figure 1, 76.6% of broad-sense heritability estimates were above 0.5, with narrow confidence intervals (supplementary table S4). Two estimators, H^2^_C_ and H^2^_O_, handling missing data differently, gave very similar estimates (supplementary table S4), both indicating that genotypic values of all cultivars were accurately estimated for most response variables. Moreover, 92.7% of the genetic coefficients of variation were above 5% and 59.1% above 20% (figure 1, supplementary table S4).

**Fig. 1.**
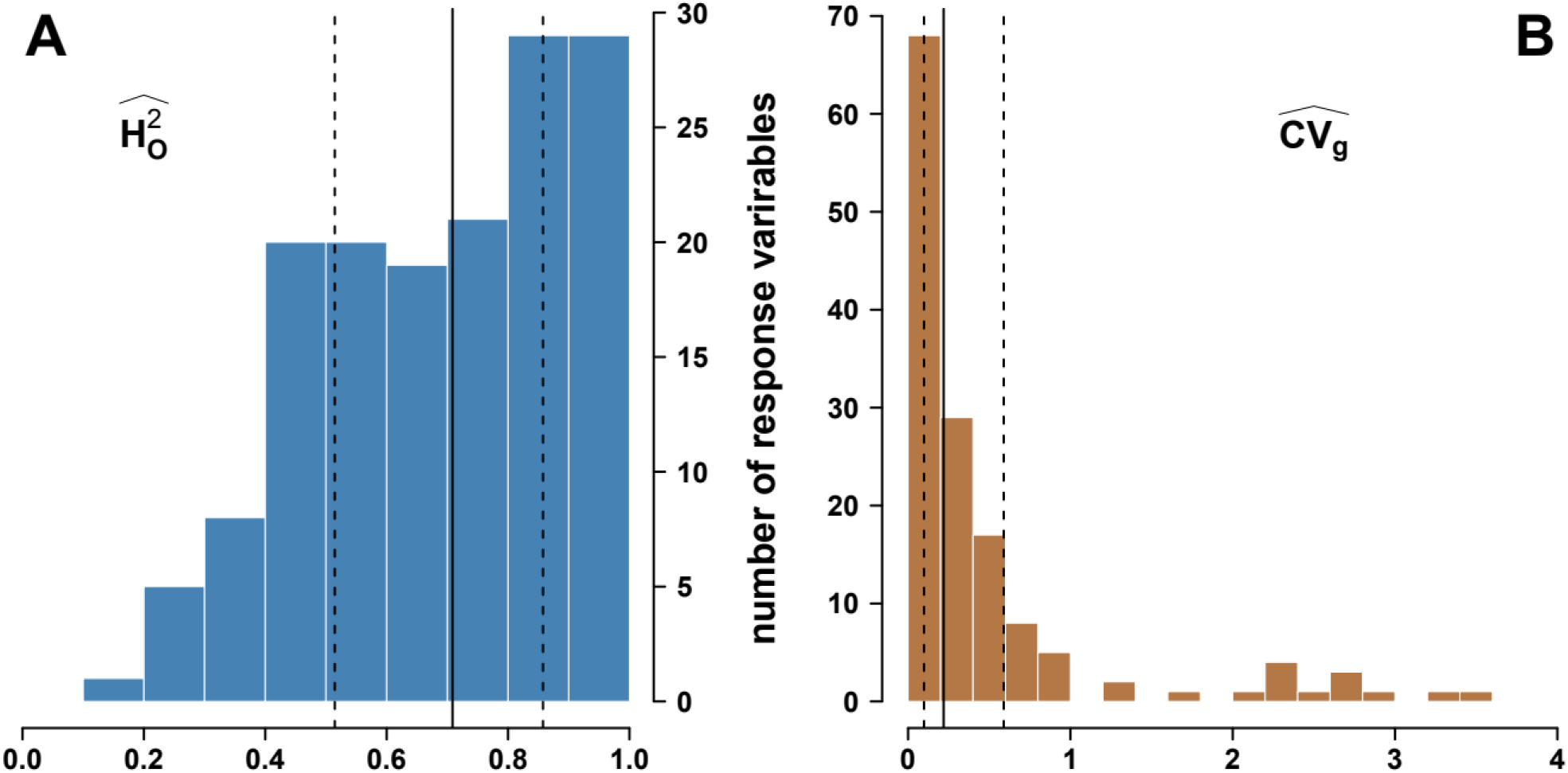
Estimation in a diverse panel of *Vitis vinifera* L. of (A) broad-sense heritabilities for 152 response variables using the estimator from Oakey et al. (2006), H^2^_O_, and (B) their genetic coefficients of variation, CV_g_. Vertical lines indicate the median (plain), and quantiles at 0.25 and 0.75 (dotted).

### Combining genotyping technologies to explain more genetic variance

We then aimed to explain the variance of these genotypic BLUPs with SNP genotypes. For that purpose, we used two sets of SNPs, the “microarray-only SNPs” (10,503 SNPs) and “microarray-GBS SNPs” (63,105 SNPs).

Because LD is known to be short in *Vitis vinifera* L. (Myles et al. 2011; Nicolas et al. 2016), we increased the SNP density initially obtained with the microarray by sequencing with complexity reduction (GBS). Raw reads had high quality along their sequences, although many displayed adapter content at their 5’ end, which had to be trimmed off. After demultiplexing, more than 95% of the reads were assigned to a cultivar. After alignment on the reference genome, the median coverage depth of regions having at least one read, averaged over cultivars, was 21.7, which allowed to accurately call both homozygous and heterozygous SNP genotypes after filtering out SNPs supported by less than 10 reads.

Compared to the microarray-only SNP set, the combined microarray-GBS set displayed a substantially higher SNP density along all chromosomes (supplementary figure S4). We then estimated the additive genetic relationships between cultivars (supplementary figure S7), confirming the weak structure in three subgroups corresponding to wine west (WW), wine east (WE) and table east (TE). The matrix of genetic relationships was used to estimate the proportion of variance in genotypic BLUPs explained by SNPs (PVE_SNPs_). Assuming an additive-only, polygenic architecture, PVE_SNPs_ was higher with microarray-GBS SNPs than with microarray-only SNPs for 97.8% of responses variables (figure 2, supplementary table S5). This showed the advantage of combining SNPs so that more QTLs are in LD with at least one genotyped SNP.

**Fig. 2.**
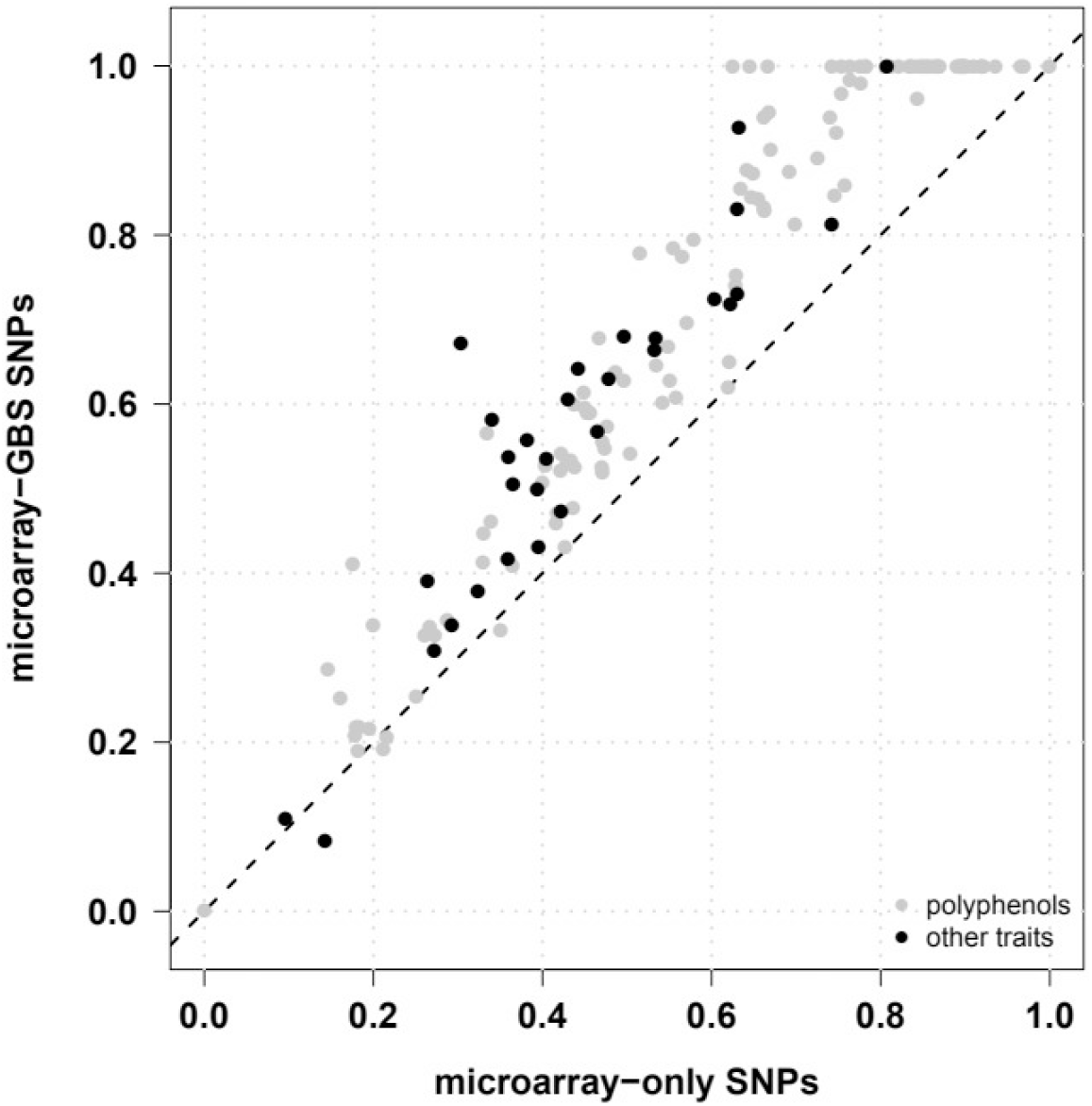
Estimation in a diverse panel of *Vitis vinifera* L. of the proportion of variance in genotypic BLUPs explained by SNPs for 152 response variables and two SNP densities, assuming an additive-only, polygenic architecture.

Models with both additive and dominance relationships either converged with difficulty or failed to, most probably because the matrix of dominance relationships was very similar to the identity matrix, making it indistinguishable from the error term (supplementary figure S7).

### QTL detection by GWAS and identification of candidate genes

The GWAS methods used in the following were first checked on two previously phenotyped traits, flower sex and berry skin color, for which the already known genetic architecture consists in a major QTL. Results were coherent with the literature (Fournier-Level et al. 2009; Picq et al. 2014): a major QTL on chromosome 2 for flower sex (around coordinate 4,769,151) and for berry skin color (around coordinate 15,753,009). Other weaker QTLs were also found, on the Unknown chromosome for flower sex (note that Tello et al. (2019) found chunks of chromosome 2 in the Unknown chromosome when building genetic maps), and on chromosome 7 and 13 for berry skin color (consistent with QTLs for skin anthocyanidin content found by Guo et al. 2015).

Each response variable phenotyped in this study was analyzed with a SNP-by-SNP model to identify significant SNPs (supplementary table S6). For each response variable, QTLs were defined as LD-based intervals around each significant SNP (supplementary table S7), and then merged when overlapping (supplementary table S8). As summarized in table 1, at least one QTL was found for 66.4% of response variables with the microarray-GBS SNPs, as compared to 57.9% with the microarray-only SNPs.

**Table 1.** Comparison between methods in terms of the number of QTLs (#QTLs) found in a diverse panel of *Vitis vinifera* L. for two SNP data sets, summed up over all response variables. Also indicated are the number of response variables with at least one QTL (#RVs), and the number of significant SNPs (#sSNPs).

| Method |  | microarray-only SNPs |  |  | microarray-GBS SNPs |  |  |
| --- | --- | --- | --- | --- | --- | --- | --- |
| Model | Software | #RVs | #sSNPs | #QTLs | #RVs | #sSNPs | #QTLs |
| SNP-by-SNP | GEMMA | 88 | 2,295 | 1,179 | 101 | 7,855 | 1,784 |
| multi-SNP | mlmm.gwas | 148 | 1,257 | 1,243 | 125 | 703 | 692 |
|  | varbvs | 118 | 266 | 257 | 119 | 258 | 257 |

To benefit from a potential gain in power, we fitted two multi-SNP models. With both, more response variables had at least one QTL compared to the SNP-by-SNP model, whatever the SNP set (table 1). Within multi-SNP methods, mlmm.gwas found more significant SNPs and QTLs than varbvs, and for more response variables. Yet, the interpretation is not straightforward as these methods do not use the same criterion for declaring a SNP as significant. Surprisingly, for mlmm.gwas, the numbers of response variables with at least one QTL, significant SNPs and QTLs werer lower with more tested SNPs.

By summing over the 150 response variables with at least one QTL, a total of 3,490 QTLs were found (supplementary table S8), which corresponded to an increase of 196% in the number of QTLs and of 70% in the number of response variables with at least one QTL, compared to applying the SNP-by-SNP method on the microarray-only SNPs. Among these QTLs, 136 were found by all three methods, while 3,001 were found by a single method only and 1,598 by multi-SNP methods only (supplementary figure S9). Furthermore, over these 150 response variables, 26 had no QTL according to the SNP-by-SNP method but at least one found by both multi-SNP methods (supplementary figure S10).

All chromosomes harbored at least one QTL (supplementary figure S11), and most QTLs found only by the multi-SNP mlmm.gwas method fell far from QTLs found by other methods (supplementary figure S12). Moreover, 90% of the QTLs found only by the SNP-by-SNP method GEMMA clustered on chromosome 2 for 64 response variables, all of them being polyphenols, in relation with the anthocyanin-related MYB genes on this chromosome (Matus et al. 2008). This is expected because GEMMA ignored LD between SNPs. In contrast, the multi-SNP varbvs method was more parsimonious, yet had enough power to identify significant SNPs in regions in which GEMMA did not identify any signal (supplementary figure S12).

To prioritize QTLs for further investigation, 489 QTLs involving 124 response variables were deemed reliable as they were found by at least two methods (supplementary table S9, supplementary figure S13). They corresponded to 59% less QTLs but 41% more response variables with at least one QTL, compared to applying the SNP-by-SNP method on the microarray-only SNPs. All chromosomes harbored at least one such reliable QTL, except chromosome 19 (figure 3).

**Fig. 3.**
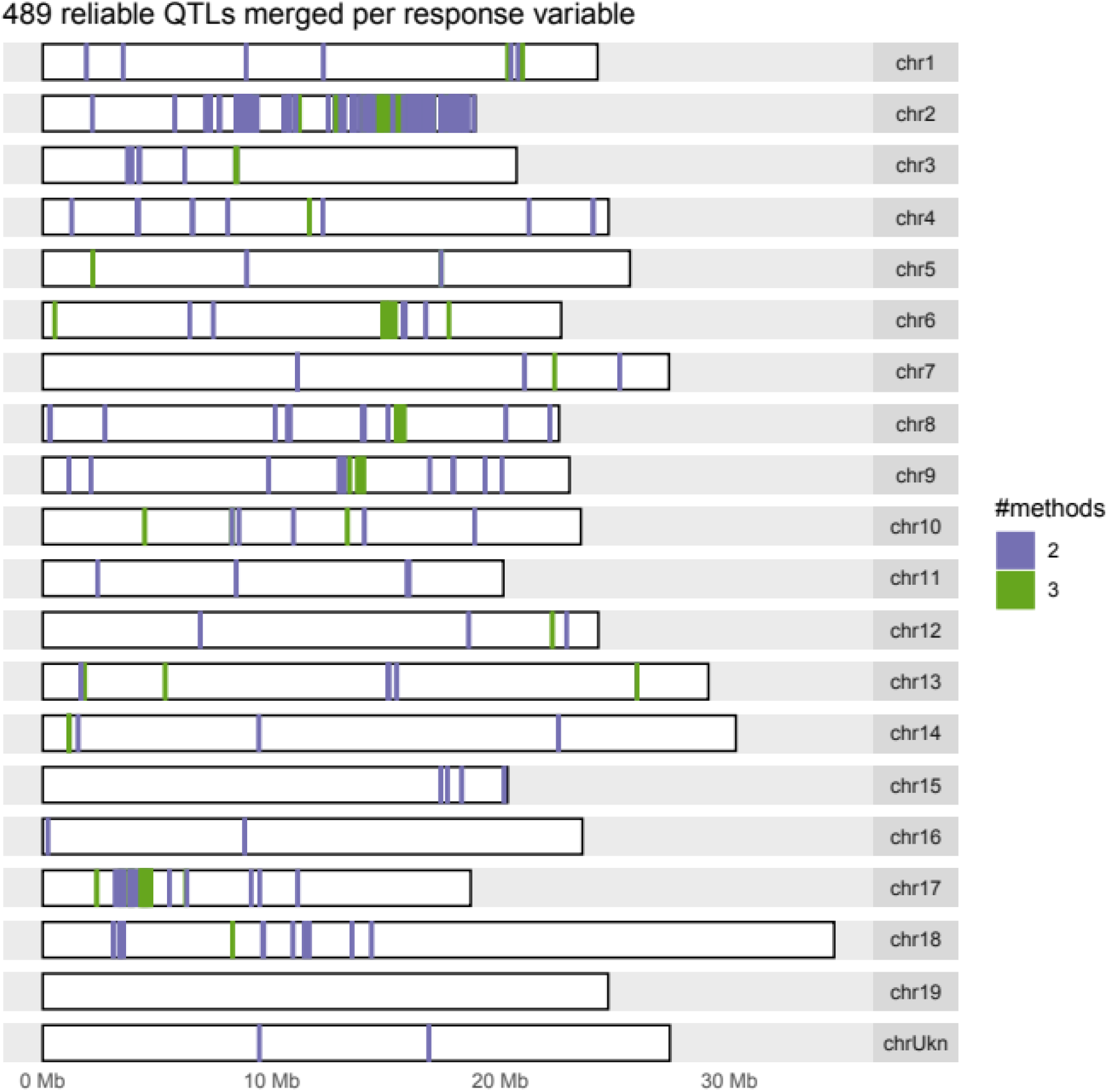
Genomic distribution of the most reliable QTLs identified by two methods in a diverse panel of *Vitis vinifera* L. after merging them over microarray-only and microarray+GBS SNP sets per response variable. The color legend indicates the number of methods that identified a given QTL.

The 489 reliable QTLs were compared with the largest list of QTLs detected in grapevine bi-parental crosses compiled so far (Vezzulli et al. 2019). Among the 22 traits in common, QTLs were found on the same chromosome for seven only (supplementary table S8): cluster number (on chromosome 7), berry weight (on chromosomes 1, 2, 8, 11, 15 and 17), malate (on chromosomes 9 and 18), glucose to fructose ratio (on chromosome 2), mean degree of polymerisation of tannins (chromosome 17), total concentration of native anthocyanins (on chromosome 2), and %B-ring methylated anthocyanins (on chromosome 2).

We also compared our reliable QTLs with significant GWAS hits published in grapevine. Only two traits (cluster and berry weight) were phenotyped in at least one other study with at least one significant GWAS hit found (Zarouri 2016; Laucou et al. 2018; Guo et al. 2019). For berry weight, out of the 10 QTLs we found, 8 were deemed new on chromosomes 1, 2, 8, 11, 15 and 17. We also found two QTLs on chromosome 8 close to a known GWAS hit (Zarouri 2016), but did not recover other hits on chromosomes 5, 17, 18 and 19 as in Zarouri (2016) and Guo et al. (2019). For cluster weight, we found two new QTLs on chromosomes 1 and 3 but did not recover other hits on chromosomes 5 and 13 (Zarouri 2016; Laucou et al. 2018).

The comparison of our reliable QTLs with the reference gene annotations detected 1926 distinct gene models (supplementary table S10). Out of these, only 980 had a proposed putative function (supplementary table S11).

### Assessment of genomic prediction and insight into genetic architectures

We assessed the accuracy of genomic prediction by cross-validation within the panel of 279 cultivars (supplementary table S12). We compared two methods assuming contrasted genetic architectures: additive infinitesimal for rrBLUP and additive sparse for varbvs. Both the median Pearson and Spearman correlation coefficients between observed and predicted genotypic values fell between 0.37 and 0.44, similarly for both SNP sets and methods (figure 4). These correlations showed substantial correlation with broad-sense heritability (supplementary figure S14), higher for varbvs (~0.65) than for rrBLUP (~0.54). However, the distributions of varbvs’ correlation coefficients were clearly multi-modal, with the majority being lower than rrBLUP’s but still a substantial fraction being higher. This confirmed the robustness of rrBLUP’s predictions irrespective of the underlying architecture (Wang et al. 2015). Yet varbvs can provide substantially better predictions than rrBLUP for traits for which the genetic architecture is likely to be rather sparse.

**Fig. 4.**
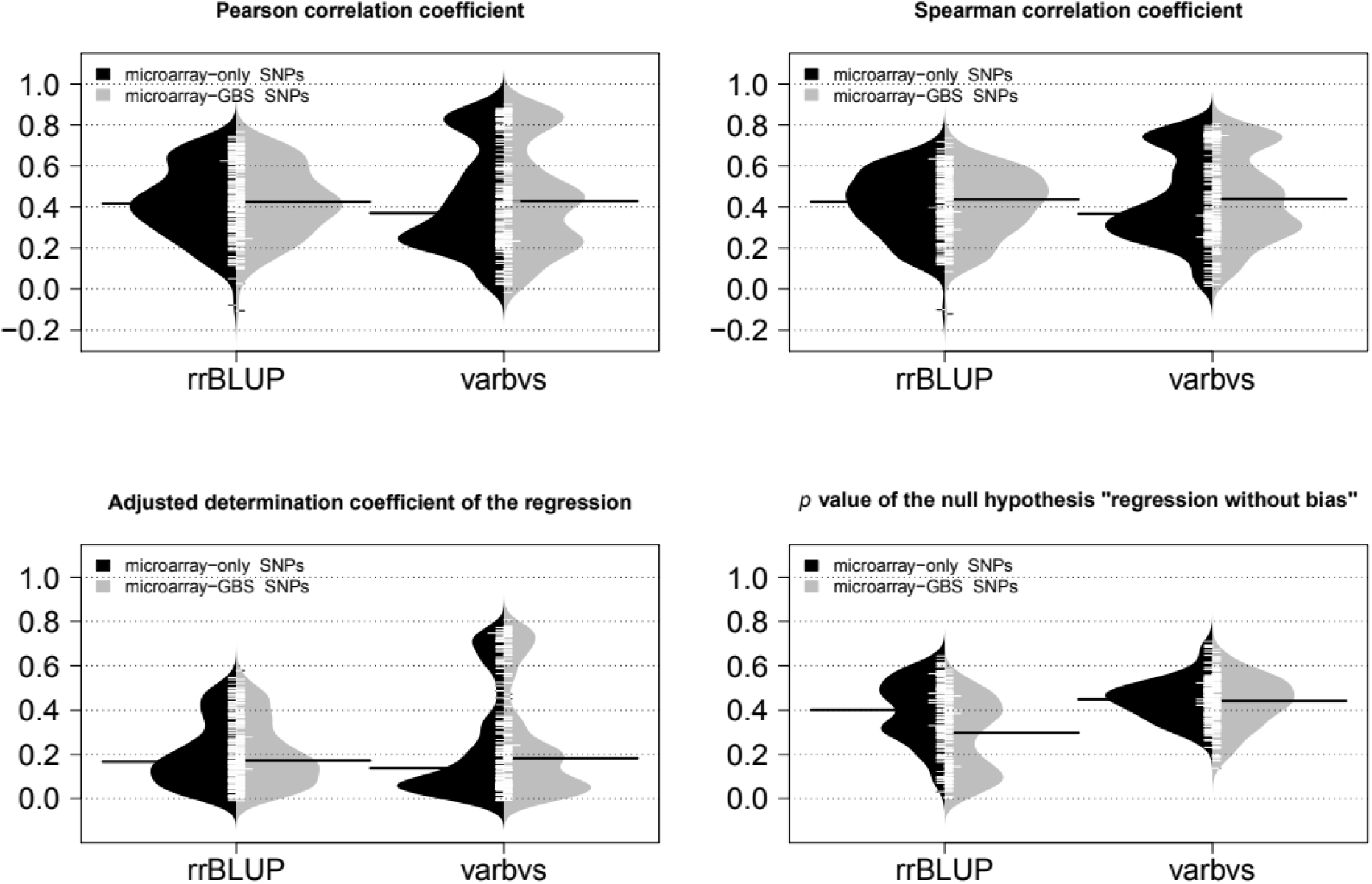
Assessment of genomic prediction accuracy within a diverse panel of *Vitis vinifera* L. by repeated 5-fold cross-validations, comparing two SNP sets (microarray-only and microarray-GBS) and two methods (rrBLUP assuming a dense genetic architecture and varbvs assuming a sparse genetic architecture) for 152 responses variables. The four displayed metrics were averaged over folds and replicates.

Moreover, rrBLUP results did not seem to depend on the SNP set whereas they were slightly better with the microarray-GBS SNPs for varbvs. This suggests that, among the extra SNPs provided by GBS, varbvs managed to identify those that improved its prediction accuracy. Concerning the *p* value of the test for no bias, varbvs showed similar values across both SNP sets, higher than rrBLUP in general and above 0.05, suggesting an absence of bias. On the contrary, rrBLUP showed lower *p* values with the microarray-GBS SNPs, suggesting that its assumption of all SNP effects being non-zero may be too strong for these traits, especially when SNP density is high.

We also used the panel of 279 cultivars as a training set to predict mean berry weight in a subset of a Syrah x Grenache progeny. With rrBLUP (respectively varbvs), this gave a Pearson correlation of 0.56 (0.35), an adjusted coefficient of regression of 0.28 (0.08), and a *p* value when testing for no bias of 1.6×10^-4^ (3.5×10^-3^). The correlation is particularly promising for rrBLUP compared to varbvs, in agreement with the results obtained by cross-validation within the panel (Pearson correlation of 0.71 with rrBLUP and 0.61 with varbvs).

Finally, combining results from both genome-wide association and genomic prediction studies provides insight into the genetic architecture of the studied traits. In table 2, trait classes are sorted according to the following metric: the difference between the accuracy of genomic prediction assuming a sparse additive genetic architecture (as implemented in varbvs) versus a dense one (as implemented in rrBLUP), using the Spearman correlation coefficient from the cross-validation above as a proxy of prediction accuracy (supplementary table S13).

**Table 2.** Types of additive genetic architecture per trait class in a diverse panel of *Vitis vinifera* L. based on the accuracy of genomic prediction assuming a sparse genetic architecture (method “varbvs) or a dense one (method “rrBLUP”) over all response variables (RVs). Also indicated are a symbol for the confidence level in the classification (+ for high, - for low), the number of reliable QTL (#relQTLs) and the broad-sense heritability estimated according to (Oakey et al. 2006) (H^2^_0_); for both, the median, quantile at 10% and quantile at 90 are given.

| Trait class | #RVs | Median of $\text{cor}_s(\text{varbvs}) - \text{cor}_s(\text{rrBLUP})$ | Additive genetic architecture | #relQTLs | $H^2_o$ |
| --- | --- | --- | --- | --- | --- |
| biochemical | 136 | +0.05 [-0.12, +0.18] | sparse (-) | 3.0 [0.0, 8.0] | 0.69 [0.41, 0.96] |
| abiotic stress | 2 | -0.04 [-0.09, +0.02] | dense (-) | 0.5 [0.1, 0.9] | 0.37 [0.21, 0.52] |
| phenological | 3 | -0.04 [-0.06, -0.03] | dense (+) | 2.0 [0.4, 2.0] | 0.80 [0.72, 0.83] |
| morphological | 5 | -0.08 [-0.08, -0.07] | dense (+) | 1.0 [0.0, 1.6] | 0.82 [0.74, 0.87] |
| agronomical | 6 | -0.12 [-0.09, +0.19] | dense (-) | 1.5 [0.5, 5.0] | 0.79 [0.38, 0.95] |

Overall, the median of this metric is positive only for response variables corresponding to biochemical traits (mostly polyphenols), suggesting a sparse genetic architecture for them. All other trait classes have a median metric which is negative, suggesting a dense genetic architecture. When taking into account the distribution of the metric, the classification in sparse or dense architecture is deemed trustworthier when the quantile interval does not include 0, which is the case for phenological and morphological traits.

Apart from the abiotic stress variable δ^13^C, all response variables had a high median broadsense heritability (around 0.7 and above), indicating higher measurement quality, hence also contributing to the trustworthiness in the suggested genetic architecture. Moreover, in the case of the biochemical response variables, the median number of reliable QTLs is higher than for the other trait classes, although there is a large variation. This is consistent with their genetic architecture deemed sparse, for which one expects to have QTLs with a large-enough effect to be found significant.

## Discussion

### Design and analysis of field trials for perennials

Acquiring phenotypic data from which genotypic values can be deduced with sufficient accuracy is a big challenge, especially because a large panel is a prerequisite to have enough power to detect QTLs (Nicolas et al. 2016). Our randomized block design certainly helped in reaching medium to high broad-sense heritability for most traits. Those with low heritability may be due to the difficulty of sampling fruits at a similar physiological stage, a particularly pressing issue for grapevine due to the strong intra- and inter-cluster heterogeneity between berries (Shahood 2017). Automatizing new protocols (Bigard et al. 2018) remains to be done to phenotype large panels.

At the first stage of the analysis, we chose to include pruning weight, the number of wooding shoots and vigour as explanatory factors in the global model, but neither flower sex nor berry color. Our rationale was that the former three are more influenced by the way the field trial is conducted than the latter two, which are more genetically determined (Fournier-Level et al. 2009; Picq et al. 2014). This approach would hence keep most genetically-based variation between genotypes for the second stage of the analysis (genome-wide association and genomic prediction). More generally, this raises the question of how to deal with multiple traits to exploit their correlations (supplementary table S14 and figure S15). Most multivariate linear models put all the traits on the same level, which complicates the understanding of their genetic architecture (Kemper et al. 2018). A more ambitious approach would leverage functional-structural plant models (Sievanen et al. 2014) but it notably requires the phenotyping of key phenological stages for the whole panel, as well as the non-destructive phenotyping of major physiological processes through time.

### Increase of genotyping density

Validating heterozygous SNP genotypes from GBS data is notoriously difficult (Swarts et al. 2014). We hence looked at the proportion of variance in BLUPs of genotypic values explained by SNP genotypes (PVE_SNPs_). The improvement obtained with the microarray-GBS set increased our trust in the genotyping and imputation procedures. Yet, PVE_SNPs_ did not equal 1 for all response variables. Several factors can underlie this discrepancy. First, empirical BLUPs of genotypic values are not fully accurate versions of the “true” genotypic values, as reflected by the broad-sense heritability. Second, the microarray-GBS SNPs may not be in strong-enough LD with all “true” QTLs. The number of required SNPs may reach half a million in grapevine (Nicolas et al. 2016), a value likely to be similar in other perennial fruit crops with low LD. Moreover, many pan-genome structural variations may remain undetected, which calls for whole-genome sequencing (Marroni et al. 2014).

### Sensitivity and specificity of QTL detection

Our study which detected many reliable QTLs benefited from a highly favorable context combining a representative panel, a proper experimental design and a large number of phenotyped traits. When comparing GWAS methods, a major misleading factor is linkage disequilibrium, which SNP-by-SNP methods do not take it into account whereas multi-SNP methods do, even though differently depending on the details of each method. We hence compared the three methods in terms of QTLs, defined here as intervals around significant SNPs, instead of significant SNPs directly. We used the genome-wide distribution of LD to define the extent of QTLs, which ignores local variations along the genome. Haplotype-based methods could provide complementary information, but is beyond the scope of this work.

We compared our reliable QTLs with those from the literature on bi-parental crosses passing a 5% genome-wide significance threshold. Therefore, when we deemed one of our QTL new, it may have been found at chromosome-wide significance threshold; nevertheless, it is reported as reliable for the first time in our study. This comparison could be achieved for a very small subset of common traits only. Part of the reason why may be that the traits studied here include an exhaustive list of polyphenols which have rarely been measured elsewhere. In addition, we faced the notorious difficulty to assess whether the same trait acronym used in different articles indeed corresponded to the same biological trait. A wider usage of a trait ontology, such as the *Vitis* ontology, seems the way forward (Krajewski et al. 2015).

Furthermore, when comparing our results on cluster and berry weights with those from the literature obtained by GWAS, we found discrepancies: several of our QTLs were new, and several QTLs reported by others were not found in our analysis. This may be due to four types of differences, (i) the composition of the association panels, (ii) the genotyping densities, (iii) the phenotyping protocols, and (iv) the statistical models. Re-analyzing these data sets was out of the scope of this work but could be done in the future depending on data availability.

### Focus on some candidate genes

For various traits, our association study identified many QTLs (supplementary tables S8 and S9) containing numerous genes (supplementary tables S10 and S11). As such, this large database is of interest *per se* to be further investigated in subsequent analyses. We chose to focus here our discussion on a subset of traits, phenolic compounds, organic acids and δ^13^C.

### Candidate genes for phenolic compounds

Our results confirm known features about the genetic regulation of phenolic compounds in grape, such as the region on chromosome 2 containing the cluster of MybA genes. It governs not only the amount and quality of anthocyanins, but also the traits concerning flavonols as already observed (Malacarne et al. 2015). Our study also confirms a large QTL for tannins composition located on chromosome 17 already detected in a Syrah x Grenache progeny (Huang et al. 2012), which contains the candidate gene *VvLAR2* (leucoanthocyanidin reductase, Vitvi17g00371). *LAR* was initially characterized as being able to catalyze the formation of catechin terminal units (Bogs et al. 2005), but more recently it was demonstrated that *VvLAR* could have an additional role in controlling the degree of polymerization (Yu et al. 2019).

Our study also identified new regions for already-studied traits, such as one involved in anthocyanin acylation and tri-hydroxylation on chromosome 13. This QTL was not detected neither in a Syrah x Pinot Noir progeny (Costantini et al. 2015), nor in a Red Globe x Muscat of Hamburg progeny (Sun et al. 2020), and is distinct from the functionally validated anthocyanin acyltransferase on chromosome 3 (Rinaldo et al. 2015 Sep 22) or the Flavonoid 3’,5’-hydroxylase cluster located on chromosome 6. This region contains two *WRKY* transcription factors (Vitvi13g00189 and Vitvi13g01916) orthologuous of *AtWRKY55* and *AtWRKY54/70 (Wang et al. 2014). WRKY* transcription factor mediates stress responses in plants (Phukan et al. 2016), and *AtWRKY70* was also described to control JA induced accumulation of anthocyanins (Li et al. 2006). In grape, anthocyanin acylation and hydroxylation are affected under abiotic stress (Ollé et al. 2011), thus those *WRKY* transcription factors appear as candidate genes to modulate anthocyanin composition.

Moreover, this is the first GWAS in grape for some phenolic compounds such as phenolic acids or dihydroflavonols. A region on chromosome 6 controlling the amount of astilbin, resulting from the rhamonsylation of taxifolin, contains four uncharacterized flavonoid O-glycosyltransferases (Vitvi06g01093, Vitvi06g01097, Vitvi06g01099, Vitvi06g01100) that could be involved in this reaction.

### Candidate genes for organic acids and δ^13^C

For citrate, no QTL had been searched for yet and our study yielded one on chromosome 3. This 56-kb region contains several candidates genes: 5 copies of allene oxide synthase (Vitvi03g00391 to 5), and the long chain acyl coA synthase 2 (Vitvi03g00388). Oxylipins formed by allene oxide synthases are precursors of jasmonates (Farmer and Goossens 2019) involved in rewiring central metabolism, thus decreasing the levels of metabolites associated with active growth like citrate (Savchenko et al. 2019). Moreover, the closest homologue of the last gene in Arabidopsis participates in oil synthesis in seed endoplasmic reticulum, where its overexpression triggers the activation of genes involved in glycolysis (Ding et al. 2020). Acyl coA synthase 2 and citrate synthase may hence compete for AcetylCoA which yields citrate when condensed with oxaloacetate.

Regarding malate, Vitvi09g00195 located on chromosome 9, possibly in a QTL found by Bayo-Canha et al. (2019) in a parental genetic map from a bi-parental progeny, encodes a chloroplastic glyoxylate/succinic semialdehyde reductase 2 which has two connections with malate synthesis. First, this enzyme may scavenge glyoxylate in the chloroplast matrix and protect photosynthesis from its adverse effects (Simpson et al. 2008). Glyoxylate is, with acetyl CoA, the direct substrate of malate synthase in glyoxysome and such diversion from the classical photorespiratory pathway was documented in Chlorella (Xie et al. 2016). Second, succinic semialdehyde dehydrogenase (SSADH) is the last enzyme in the gamma-aminobutyric acid shunt of the TCA cycle (Zarei et al. 2017), forming succinate that is readily oxidized to fumarate, the precursor of malate in mitochondria. In another new malate QTL on chromosome 12, Vitvi12g00505 encodes a cytosolic aconitate hydratase that may complement the activities of the mitochondrial and glyoxysomal ones, involved in the metabolism of dicarboxylate and glyoxylate, respectively. On chromosome 18, Vitvi18g01038 encodes V-type proton ATPase subunit a2, a part of the hydrophobic V0 rotor that generates the membrane potential essential for the storage of organic acids in grapevine fruit (Terrier et al. 2001). Noticeably, in a Riesling x Gewurztraminer progeny, subunits G of V-ATPase on chromosomes 8 and 13 were suggested as candidate genes for acidity QTLs (Duchêne et al. 2020).

Relevant candidate genes were also found under novel QTLs for δ^13^C, in particular Vitvi08g02203 on chromosome 8 which encodes the transcriptional regulator *TAC1*-like. In rice it corresponds to a major QTL controlling tiller angle, with a direct influence on leaf exposition and shading (Yu et al. 2007). We also noticed the presence of different candidate genes involved in stele expansion or differentiation, like *CASP-*like proteins and Lonesome Highway (*LHW*) transcription factor.

### Genomic prediction, and the wider goal of understanding genetic architectures

The accuracy of genomic prediction, assessed for the first time for such a large number of traits by cross-validation within a grapevine diversity panel, reached promising levels according to the median Pearson correlation (around 0.4) even though the coefficient of determination remains substantially lower (around 0.17, figure 4). Nevertheless, in breeding, one mostly aims at accurately predicting the ranks of candidate genotypes, and the median Spearman correlation around 0.4 is relevant for that purpose.

Cross-validation results are interesting *per se* as they provide an upper threshold on prediction accuracy. Yet in breeding, the ultimate goal lies in training a model on a reference panel to predict genotypic BLUPs in a segregating population. When testing this with a subset of a Syrah x Grenache progeny not belonging to the panel, the accuracy metrics were lower than with within-panel cross-validation, yet they displayed the same trend in terms of methods. Ridge regression model (rrBLUP) performed better than the sparse regression model (varbvs), which may be due to the essentially infinitesimal architecture of the trait despite a few larger QTL segregating for this trait in the progeny (Doligez et al. 2013). This promising result was studied in more details with other traits and other progenies in another work (Brault et al. 2021), as was done in other perennial fruit crops (Minamikawa et al. 2017; Roth et al. 2020).

In terms of genetic architectures, we focused on additive ones and attempted at distinguishing trait classes with a sparse vs. dense architecture. Leaving aside the trait class “abiotic stress” which had a low broad-sense heritability, our results based on prediction accuracy indicated a sparser architecture for biochemical traits, versus a denser one for phenological, morphological and agronomical traits. In the framework of genotype-phenotype maps, this may correspond to the fact that biochemical traits are closer, in a causal sense, to genetic variation (such traits are sometimes called “endophenotypes”), hence making QTL detection easier. On the opposite, the other trait classes are more integrated, in the sense of resulting from multiple developmental and ecophysiological processes (Granier and Vile 2014). Moreover, the determination of genetic architecture is also known to depend on sample size and LD extent (Wimmer et al. 2013). In contrast to expectations on annual plant breeding populations, we identified traits with better prediction accuracy assuming a sparse architecture rather than a dense one, in spite of the rather small sample size of our panel. This was likely due to the short LD within this diversity panel, a notable feature of perennial plants, although these results may not stand for grapevine bi-parental breeding populations with longer LD.

## Conclusion

This work demonstrated the feasibility of performing a genome-wide association study in a perennial fruit crop such as grapevine for numerous, mostly complex traits related to various aspects of plant biology and breeding. A key ingredient for field trials remains the experimental design necessary to achieve a high broad-sense heritability. We also provided dense genotyping data for further studies on the panel, although, given the low LD, an even higher number of SNPs would be advantageous. In terms of GWAS, we confirmed that a gain in power is possible when using multiple-SNP models. Overall, we identified new QTLs as well as promising genes under them, leading to mechanistic hypotheses yet to be tested. In terms of genomic prediction, we provided a distribution of prediction accuracy across many traits likely to have various genetic architectures. We confirmed the usage of the RRBLUP/GBLUP model assuming a dense architecture as a relevant default. Yet we showed that a model assuming a sparse architecture can reach higher prediction accuracy for some traits, notably in the case of traits closer to the genetic variation. As such, our work provided important results for the contribution of genomic prediction in breeding perennial crop species.

## Author contribution statement

PT, AD, JMB and LLC initiated the project. AD and JPP conceived the experimental design in the field. GB and YB installed and managed the field trial under the supervision of JPP and LLC. GB, YB, JPP, AD, LLC, RB, TL, JMB, VL, PT collected phenotypic data on clusters and berries in 2010-2012. CR and LLC collected organic acid data from berries in 2011-2012. LLC, VC and JPP conceived the experimental design in 2014-2015. AF, GB, YB and LLC collected phenotyping data in 2014-2015 and extracted DNA samples for the first GBS phase. IB tested the presence of viruses. MR, GB, YB and LLC prepared samples before polyphenols, β-damascenone and pDMS extraction. VB collected β-damascenone and pDMS data. LLC and TF conceived the experimental design for the GBS. AL extracted DNA samples for the second GBS phase and made the libraries. TF wrote all the code and performed the statistical analyses. TF, AD and CR interpreted the results. CR and NT analyzed candidate genes. TF drafted the manuscript. All authors contributed critical revision of the work and approved the manuscript.

## Ethical standards

The authors declare that the experiments comply with the current laws of the country in which they were carried out.

## Data availability

The data that support the findings of this study are openly available, for sequences at the NCBI as BioProject PRJNA489354 and, for all the rest, at the Data INRAE repository at *URL which will be made public upon publication*.

## Code availability

The code that supports the findings of this study is openly available at the Data INRAE repository at *URL which will be made public upon publication*. Many of the custom functions we used are available in a R package for reproducibility purposes, rutilstimflutre (Flutre 2019).

## Acknowledgments

We thank all the agents from the grapevine germplasm collection maintained at INRAE Vassal, Valérie Miralles and Jean-François Ballester from AGAP-PPB for phenotyping of organic acids, Pierre Mournet from AGAP-GPTR and the GenoToul platform for genotyping by sequencing, and Bertrand Pitollat and the South Green platform for computing.

## Footnotes

Supplemental material available at the Data INRAE repository at *URL which will be made public upon publication*.

## Figures and tables

As this is a first submission (i.e., not a revised manuscript), we chose to included the figures and tables within the main text above.

